# Posttranscriptional tuning of gene expression over a large dynamic range in synthetic tobacco chloroplast operons

**DOI:** 10.1101/2024.01.03.574089

**Authors:** Qiguo Yu, Tarinee Tungsuchat-Huang, Alexander Ioannou, Alice Barkan, Pal Maliga

**Affiliations:** Waksman Institute of Microbiology, Rutgers University, Piscataway, NJ 08854, USA; Institute of Molecular Biology, University of Oregon, Eugene, OR 97403, USA; Department of Plant Biology, Rutgers University, New Brunswick, NJ 08901, USA

**Keywords:** posttranscriptional regulation, chloroplast gene expression, synthetic operons, PPR10 RNA binding protein, tRNA *cis*-element

## Abstract

Achieving balanced gene expression within synthetic operons requires a spectrum of expression levels. Here we investigate the expression of *gfp* reporter gene in tobacco chloroplasts, guided by variants of the plastid *atpH* 5’ UTR, which harbors a binding site for PPR10, a protein that activates *atpH* at the post-transcriptional level. Our findings reveal that endogenous tobacco PPR10 confers distinct levels of reporter activation when coupled with the tobacco and maize *atpH* 5’ UTRs in different design contexts. Notably, high GFP expression was not coupled to stabilization of monocistronic *gfp* transcripts in dicistronic reporter lines, adding to the evidence that PPR10 activates translation via a mechanism that is independent of its stabilization of monocistronic transcripts. Furthermore, the incorporation of a tRNA upstream of the UTR nearly abolishes *gfp* mRNA (and GFP protein), resulting in a substantial reduction in GFP accumulation. When combined with a mutant *atpH* 5’ UTR, the tRNA leads to an exceptionally low level of transgene expression. Collectively, this approach allows for tuning reporter gene expression across a wide range, spanning from 0.02% to 25% of the total soluble cellular protein (TSP). These findings highlight the toolbox available for plastid synthetic biology applications requiring multigene expression at varying levels.

## INTRODUCTION

Plastid genomes are appealing targets for the expression of foreign products, with advantages that include the relative ease of manipulating their genomes, the bio-containment of foreign products, and a compartment that provides a variety of metabolites and reducing power (Scharff and Bock, 2014). In particular, their prokaryotic ancestry endowed plastids with polycistronic transcription units, which facilitates the stacking of transgenes into operons that can be leveraged for metabolic engineering (Boehm and Bock, 2018). Indeed, several metabolites have been produced by transferring entire biosynthetic pathways into plastids (Fuentes et al., 2018; Jensen and Scharff, 2019). Additionally, transferring the carbon-concentrating mechanism (CCM) from cyanobacterium into plastids to improve photosynthesis illustrates ways in which agronomic traits can be modified by plastid engineering (Lin et al., 2014; Long et al., 2018; Chen et al., 2023).

Fulfillment of this promise requires control over the expression level of each plastid transgene such that proteins are produced in optimal stoichiometries. However, the repertoire of regulatory sequences used to tune plastid transgene expression remains largely dependent on a handful of promoters and 5’ and 3’ UTRs (Boehm and Bock, 2018), including both native chloroplast elements and elements derived from bacteria or bacteriophage (Occhialini et al., 2019). Some attempts at metabolic engineering used complete bacterial operons including intergenic sequences, whereas others incorporated the chloroplast-derived “intercistronic expression element (IEE) to increase the expression of internal genes. Although high expression levels of specific proteins have been achieved in this way, relative expression levels within synthetic chloroplast operons have been unpredictable and variable (Lossl et al., 2005; Hasunuma et al., 2008; Krichevsky et al., 2010; Bohmert-Tatarev et al., 2011; Harada et al., 2014).

Recent advances in understanding the mechanisms of chloroplast gene expression provide new avenues to address this challenge. In particular, nucleus-encoded helical repeat RNA binding proteins play a prominent role in stabilizing specific chloroplast RNAs and activating their translation (Barkan and Small, 2014; Hammani et al., 2014). The pentatricopeptide repeat (PPR) proteins are the largest and best understood family of this type. PPR proteins bind specific RNA sequences via modular one repeat: one nucleotide interactions, with nucleotide specificity determined in part by a predictable amino acid code (Barkan et al., 2012; Shen et al., 2016). Maize PPR10 serves as a paradigm for the mode of action of a subset of PPR proteins. PPR10’s best characterized *cis*-element maps in the *atpH* 5’ UTR. The binding of PPR10 to the *atpH* site dramatically increases *atpH* expression by blocking 5’-to-3’ RNA degradation and by increasing translational efficiency (Pfalz et al., 2009; Prikryl et al., 2011; Rojas et al., 2018). The ability of the IEE to boost the expression of operon-internal genes (Zhou et al., 2007) results from an analogous mechanism: the IEE binds the protein HCF107, which activates *psbH* translation and blocks the exonucleolytic degradation of RNA adjacent to its binding site; (Sane et al., 2005; Hammani et al., 2012).

Application of these principles to chloroplast synthetic biology is just beginning. For example, several PPR binding sites have been tested in transplastomic constructs to investigate their potential as substitutes for the classic IEE. The binding sites of HCF152 upstream of *petB,* and CRR2 upstream of *ndhB* showed potential as alternative activating sequences (Legen et al., 2018). In addition, we demonstrated that the regulated expression of nucleus-encoded PPR10 variants with modified sequence specificity regulates the expression of chloroplast transgenes with cognate *atpH* 5’UTR sequences (Rojas et al., 2019).

The objective of this study was to devise approaches to tune protein output from cotranscribed genes by modifying intercistronic sequences containing variants of the PPR10 binding site. We also examined a tRNA *cis*-element as a means to further alter transgene expression levels. We report here that these *cis*-elements can be tailored to tune the expression of a plastid GFP reporter across a large dynamic range, producing GFP between 0.02% to 25% of total soluble cellular protein. Our findings show that native plastid *cis*-elements can be tailored to tune plastid transgene expression across a large dynamic range in synthetic operons.

## RESULTS

### Reporter constructs

We evaluated the effects of *atpH* 5’ UTR variants on the expression of a GFP reporter in both monocistronic (**Fig. 1B**) and dicistronic contexts (**Fig. 1C**). These constructs were similar in design to those we used previously to assess the effects of nucleus-encoded PPR10 variants and to test the rules of expressing polycistronic mRNAs in chloroplasts (Rojas et al., 2019; Yu et al., 2019) The constructs placed the 97-nucleotides directly upstream of the tobacco or maize *atpH* start codon (**L*atpH*^Nt^ and L*atpH*^Zm^, respectively,** **Fig. 1A**) adjacent to the GFP open reading frame, with additional features added upstream of L*atpH* in some cases (**Fig. 1C**). The *atpH* 5’ UTRs from tobacco and maize show one nucleotide mismatch within their minimal PPR10 binding sites (BS), one extra nucleotide in the spacing region between the maize BS and Shine-Dalgarno sequence, and numerous mismatches in the AT-rich sequence upstream of the BS (**Fig. 1A**). Some experiments included derivatives of maize L*atpH^Zm^*in which the seventh and eighth nucleotides of the minimal PPR10 binding site were changed to either GG (L*atpH*^GG^) or AA (L*atpH*^AA^) (**Fig. 1A**). These mutations drastically reduce the affinity for maize PPR10 *in vitro* (Barkan et al., 2012) and prevent activation by native tobacco PPR10 *in vivo* (Rojas et al., 2019; Yu et al., 2019).

**Figure 1.**
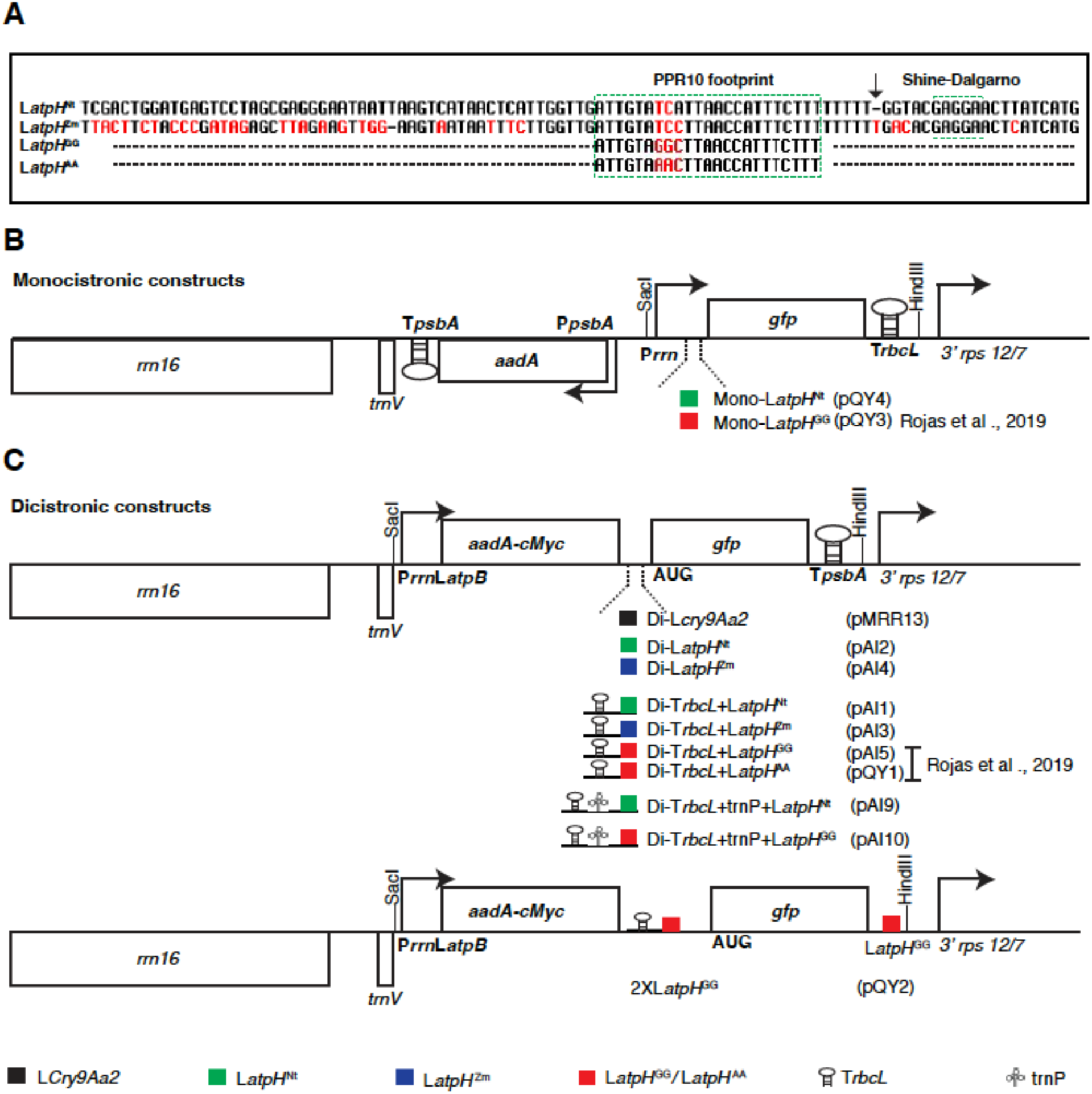
Plastid transformation vectors. **A)** Alignment of the 97 nucleotides upstream of the tobacco (Nt) and maize (Zm) plastid *atpH* translation initiation codon (AUG), containing the wild type PPR10 binding sites and two maize binding site variants (“GG” and “AA”). The PPR10 footprint (Prikryl et al., 2011) and predicted Shine-Dalgarno sequence are boxed. **B)** Monocistronic pPRV111A vector derivatives (Zoubenko et al., 1994). The *gfp* gene is transcribed from the ribosomal RNA operon promoter (P*rrn*) fused with *atpH* 5’ UTR variants (shown below the map). Construct names are in parentheses to the right. The *aadA* spectinomycin resistance gene is in a *psbA* promoter-3’ UTR (P*psbA*/*TpsbA*) cassette. The *rbcL* and *psbA* 3’ UTRs (T, terminators) are illustrated as stem-loops (T*rbcL* and T*psbA*). *rrn16*, *trnV* and 3’*rps12/7* are plastid genes that provide substrates for homologous recombination with the plastid genome. **C)** Dicistronic transformation vectors carrying the *aadA* and *gfp* genes. The *aadA* gene is expressed from the P*rrn* promoter fused with the *atpB* gene leader (L*atpB*). The intergenic regions assayed here are listed below the map. Construct names are in parentheses to the right.

The monocistronic constructs contained either the endogenous tobacco *atpH* 5’ UTR (**Mono-L*atpH*^Nt^, pQY4,** **Fig. 1B**), or, for comparison, the L*atpH*^GG^ variant of the maize *atpH* 5’UTR, which does not bind the wild-type PPR10 due to mutations in the binding site (Rojas et al., 2019) (**pQY3,** **Fig. 1B**). The dicistronic reporter constructs mimicked the native environment of PPR10 binding sites in intergenic regions of cotranscribed genes, by placing variants of the *atpH* 5’ UTR between cotranscribed *aadA* and *gfp* genes (**Fig. 1C**). The intercistronic region of the first family of constructs harbors wild-type L*atpH*^Nt^ and L*atpH*^Zm^ (**pAI2 and pAI4,** **Fig. 1C**). A second set of constructs includes a portion of the *rbcL* 3’ UTR (T*rbcL*) upstream of different L*atpH* variants; T*rbcL* forms a stem-loop structure that protects the mRNA from degradation by 3’ to 5’ exonucleases, acting as an mRNA stabilizing element (Stern and Gruissem, 1987). The variant *cis*-elements include the wild-type tobacco or maize *atpH* 5’ UTR, and two variants of the maize sequence, L*atpH*^GG^ and L*atpH*^AA^ shown previously to disrupt binding by tobacco PPR10 *in vivo* (Rojas et al., 2019). The third family has two members, Di-T*rbcL*+trnP+L*atpH*^Nt^ and Di-T*rbcL*+trnP+L*atpH*^GG^ (**pAI9 and pAI10,** **Fig. 1C**) in which T*rbcL* and L*atpH*^Nt^ are separated by the plastid *trnP* gene of *Medicago truncatula* (AC093544). The tRNA *cis*-element is projected to trigger efficient cleavage by tRNA processing enzymes, which might thereby promote the formation of monocistronic *gfp* mRNA (Chakrabarti et al., 2006; LaManna et al., 2023). The last set of dicistronic family is represented by a single construct, where the T*psbA* in Di-T*rbcL*+L*atpH*^GG^ (pAI5) was replaced with L*atpH*^GG^ (**Fig. 1C, lower panel, pQY2**). The *gfp* transcript from pQY2 tobacco was expected to be unstable due to the lack of mRNA protection from degradation in the 3’ end. A construct employing the leader sequence of the *Bacillus thuringiensis cry9Aa2* gene served as a control; L*cry9Aa2* was previously shown to yield only stable dicistronic mRNA(**Fig. 1C, pMRR13**) (Yu et al., 2017; Yu et al., 2020). We previously reported results in tobacco for the monocistronic and dicistronic constructs harboring the maize L*atpH*^GG^ UTR (pQY3 and pAI5, respectively) (Rojas et al., 2019) but they were analyzed again here as a point of comparison for other constructs.

Plastid transformation was carried out by the biolistic process, followed by selection for spectinomycin resistance encoded in the *aadA* gene. The constructs integrated into the plastid genome between the *trnV* and *3’-rps12/7* operons, a location where there is no readthrough transcription from the *rrn* operon (Zoubenko et al., 1994). Uniform transformation of plastid genomes was confirmed by DNA gel blot analyses (**Supplemental Fig. S1**). One line from each construct was selected for in-depth analysis.

### The endogenous tobacco *atpH* 5’ UTR yields robust reporter expression in a monocistronic context

Our recent work has demonstrated the efficacy of combining engineered PPR10 protein (PPR10^GG^) with its cognate binding site (“GG site”) to efficiently activate plastid transgene expression (Rojas et al., 2019; Yu et al., 2019). Here, we first sought to assess the efficacy of the endogenous tobacco *atpH* 5’ UTR itself to activate reporter expression in a simple monocistronic context by evaluating GFP accumulation from the Mono-L*atpH*^Nt^ line (**pQY4,** **Fig. 1B**, **Fig. 2**). For reference, we included the maize Mono-L*atpH*^GG^ line (**pQY3,** **Fig. 1B**), which was shown to yield low level GFP expression in tobacco chloroplasts (Rojas et al., 2019). GFP was easily visible as a band in Coomassie stained protein gels from the leaf extract of the Mono-L*atpH*^Nt^ plant (**upper panel in** **Fig. 2A**). Protein quantification indicates it accumulated about 16% of the total soluble protein (TSP) (**Fig. 2A**, **Table 1**). Based on extrapolation of a protein dilution series of the Mono-L*atpH*^Nt^ line and western blot, we estimate GFP comprises 1% of TSP from the Mono-L*atpH*^GG^ line (**lower panel in** **Fig. 2A**, **Table 1**), which is consistent with previous results (Rojas et al., 2019). Differences in GFP accumulation levels are also evident upon illumination with UV light (**Fig. 2B**). The Mono-L*atpH*^Nt^ line displayed a growth defect, in part, due to metabolic burden imposed by the high levels of GFP expression and, in part, to the additional copies of L*atpH*^Nt^ that compete with native PPR10 binding endogenous *atpH* 5’ UTR (**Supplemental Fig. 2**). These results show that the endogenous *atpH* 5’ UTR can be exploited to achieve high-level transgene expression and the Mono-L*atpH*^GG^ signals for a moderate level transgene expression.

**Figure 2.**
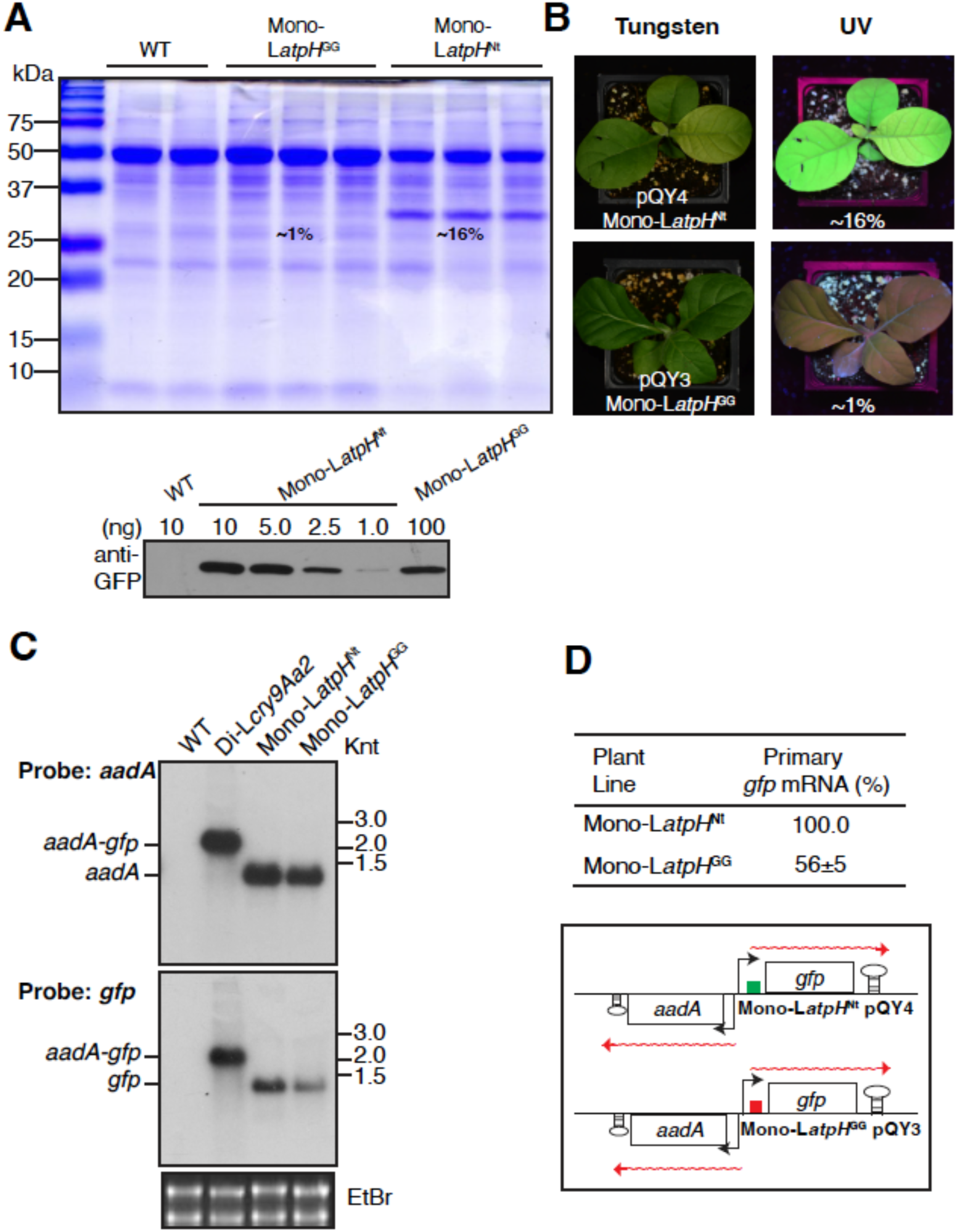
The tobacco *atpH* 5’ UTR in the monocistronic construct confers high-level GFP expression. **A)** Coomassie Brilliant Blue R250 stained SDS gel (upper panel; 5μg of total soluble fraction of leaf lysates per lane) and immunoblot analysis (lower panel) to compare the GFP abundance between Mono-*atpH*^Nt^ and Mono-*atpH*^GG^. The intensity of visible GFP bands from Mono-*atpH*^Nt^ samples was quantified with 1D-Multi Lane Densitometry (Alphaimager 2200) based on three biological replicates (mean±SD, n=3). Fold differences between Mono-*atpH*^Nt^ and Mono-*atpH*^GG^ were then quantified on western blots in ImageJ (version 2.0.0-rc-38/1.50b) using Mono-L*atpH*^Nt^ (∼16% of TSP) leaf extracts as a reference. The amount (ng) of total soluble protein per lane is indicated above the line (lower panel). **B)** Detection of GFP in leaves by illumination with UV light (UVL-56 Handheld Light (UVP Corporation, Upland, CA). **C)** RNA gel blot hybridizations showing *aadA* and *gfp* mRNA accumulation in transplastomic plants harboring the L*Cry9Aa2*, Mono-*atpH*^Nt^ and Mono-*atpH*^GG^ constructs. The ethidium bromide-stained gel below the blot shows rRNA abundance as a loading control. Knt, Kilonucleotide. **D)** Quantification of *gfp* mRNA in plants transformed with the Mono-*atpH*^Nt^ and Mono-*atpH*^GG^ vectors based on three biological replicates (mean±SD, n=3). RNA blots in Fig. 2c were imaged with a Typhoon RGB scanner (GE LifeSciences) and quantified with ImageQuant software. The abundance of mRNA was calculated relative to Mono-L*atpH*^Nt^ (100%).

**Table 1.** Levels of GFP accumulation in the leaves of transplastomic plants based on three biological repeats.

| Plant lines | Promoter+TCR1 | Intergenic region | GFP (%):<br>Average+/-SD | Terminator |
| --- | --- | --- | --- | --- |
| pAI2 | Nt-*Prrn*+*LatpB* | *LatpH*<sup>Nt</sup> | 17±2 | *TpsbA* |
| pAI4 | Nt-*Prrn*+*LatpB* | *LatpH*<sup>Zm</sup> | 25±3 | *TpsbA* |
| pAI1 | Nt-*Prrn*+*LatpB* | *TrbcL*+ *LatpH*<sup>Nt</sup> | 13±1 | *TpsbA* |
| pAI3 | Nt-*Prrn*+*LatpB* | *TrbcL*+*LatpH*<sup>Zm</sup> | 20±2 | *TpsbA* |
| pAI5 | Nt-*Prrn*+*LatpB* | *TrbcL*+*LatpH*<sup>GG</sup> | 1.8±0.1 | *TpsbA* |
| pQY1 | Nt-*Prrn*+*LatpB* | *TrbcL*+*LatpH*<sup>AA</sup> | 3.3±0.7 | *TpsbA* |
| pAI9 | Nt-*Prrn*+*LatpB* | *TrbcL*+tRNA+*LatpH*<sup>Nt</sup> | 0.4±0.1 | *TpsbA* |
| pAI10 | Nt-*Prrn*+*LatpB* | *TrbcL*+tRNA+*LatpH*<sup>GG</sup> | 0.02±0.01 | *TpsbA* |
| pQY2 | Nt-*Prrn*+*LatpB* | *TrbcL*+*LatpH*<sup>GG</sup> | 0.2±0.1 | *LatpH*<sup>GG</sup> |
| pQY4 | Nt-*Prrn*+*LatpH*<sup>Nt</sup> | None: monocistronic construct | 16±1 | *TrbcL* |
| pQY3 | Nt-*Prrn*+*LatpH*<sup>GG</sup> | None: monocistronic construct | 1±0.1 | *TrbcL* |

RNA gel blots (**Fig. 2C, 2D**) showed that the Mono-L*atpH*^Nt^ line yielded approximately 2-fold more monocistronic mRNAs than Mono-L*atpH*^GG^, implying a positive role of the endogenous *atpH* 5’ UTR in mRNA stabilization. The fact that GFP abundance was ∼16 fold higher in L*atpH*^Nt^ than in L*atpH*^GG^, whereas GFP mRNA was only 2-fold higher demonstrates that PPR10 binding enhances translation via the *atpH* 5’ UTR even in the context of monocistronic mRNA. This is consistent with prior work showing that PPR10’s activation of *atpH* occurs largely *via* its effect on translational efficiency (Zoschke et al., 2013).

### The *atpH* 5’ UTR functions as an efficient intercistronic expression element

The original intercistronic expression element (IEE), which was derived from the *psbH* 5’UTR (including the HCF107 binding site), has proven to be effective at enhancing the expression of a wide variety of plastid transgenes (Zhou et al., 2007). We set out to test a portion of the *atpH* 5’ UTR (including the PPR10 binding site) for utility as an intercistronic expression element. The dicistronic constructs discussed in this section differ only with respect to sequences between the *aadA* and *gfp* genes (**Fig. 1C, upper panel**). The goal was to determine how each variant intergenic region affects GFP and mRNA abundance.

The first group comprises constructs with the wild type tobacco or maize *atpH* 5’ UTR in the intergenic region (Di-L*atpH*^Nt^ and Di-L*atpH*^Zm^, respectively). RNA gel blot hybridizations showed that dicistronic mRNAs are the dominant form of transcripts (**Fig. 3A**). The *aadA* and *gfp* probes detect some monocistronic mRNA on blots of Di-L*atpH*^Nt^ and Di-L*atpH*^Zm^ plants, likely stabilized by PPR10 binding (**Fig. 3A**). The control, the *cry9Aa2* 5’ UTR in the *aadA*-*gfp* intergenic region, has been shown previously not to yield processed monocistronic RNA in plastids (Yu et al., 2020). The monocistronic *gfp* mRNA with the wild type tobacco *atpH* 5’ UTR is ∼4 times more abundant, indicating more efficient protection of the mRNA by the cognate wild type tobacco PPR10 protein (**Di-L*atpH^Nt^*,** **Fig. 3A**, **Fig. 3B**, **Table 1**), compared to the maize Di-L*atpH^Zm^* 5’ UTR that contains a single mismatch in the 17 nt binding site (**Fig. 1A**). Either the native tobacco PPR10 protein discriminates between the maize and tobacco binding sites and more efficiently protects the tobacco variant or the sequence upstream of the PPR10 binding site in maize provides more ready access to initiating endonucleolytic cleavages than the tobacco sequence.

**Figure 3.**
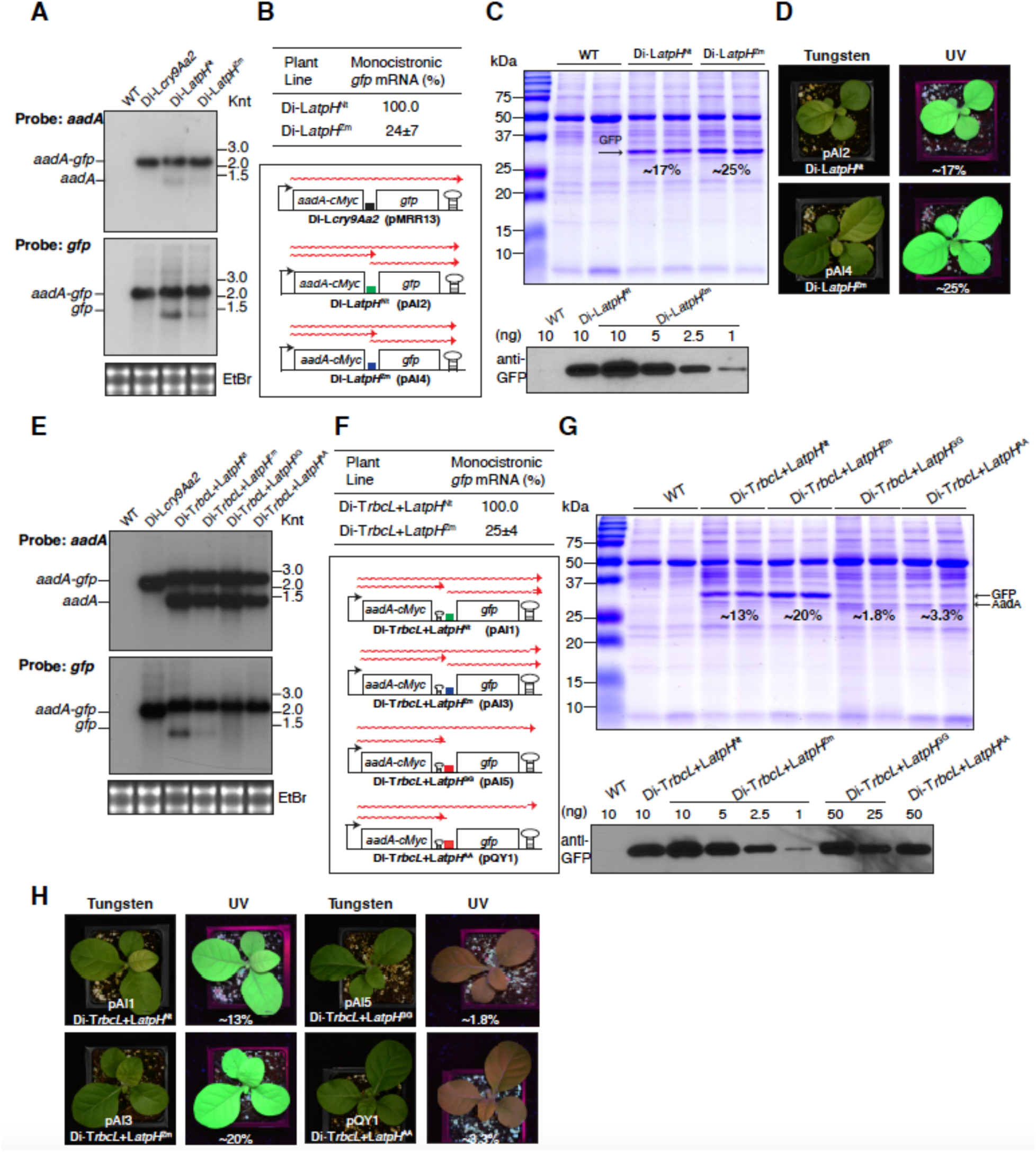
The impact of *atpH* 5’ UTRs on mRNA processing, stabilization and translational activation in dicistronic operons. **A)** RNA gel blot analysis using a probe for *aadA* (upper panel) and *gfp* (lower panel). Each lane was loaded 2μg total leaf RNA. Knt, Kilonucleotide. The ethidium bromide-stained gels showing the 28S and 18S rRNAs below the blot are loading controls. Knt, Kilonucleotide. **B, F)** Quantification of monocistronic *gfp* mRNA, as described in Fig. 2D. Below is a schematic representation of the dicistronic operon with primary transcripts and processed mRNAs. **C, G)** Quantification of GFP in Coomassie Brilliant Blue R250 stained Urea gel (upper panel) and immunoblot analyses (lower panel). Note Fold differences among Di-T*rbcL*-L*atpH*^GG^ and Di-T*rbcL*-L*atpH*^AA^ were quantified on western blots in ImageJ (version 2.0.0-rc-38/1.50b) using Di-T*rbcL*-L*atpH*^Zm^ (∼20% of TSP) leaf extracts as a reference. Note the urea alters the relative mobility of GFP (predicted ∼29kDa) and AadA(predicted ∼31kDa). **D, H)** GFP in the leaves was visualized by fluorescence under 366 nM UV light upon illumination with a UVL-56 Handheld Light (UVP Corporation, Upland, CA). **E)** RNA gel blot analysis detects dicistronic and processed monocistronic mRNAs. Knt, Kilonucleotide.

Unanticipatedly, the GFP accumulation in the leaves of tobacco plants was lower in Di-L*atpH*^Nt^ (17% of TSP) than in Di-L*atpH*^Zm^ (25% TSP) (**Fig. 3C**). Efficient translation of the internal (second) ORF is expected, as translation initiates independently from different ribosome binding sites within the same polycistronic mRNA (Zoschke et al., 2013; Chotewutmontri and Barkan, 2016; Yu et al., 2020). Having more GFP programmed from the maize L*atpH*^Zm^ mRNA is surprising given the expectation that tobacco PPR10 is predicted to bind with higher affinity to its native binding site. Possible explanations for this unanticipated observation are discussed below. In any case, the very high GFP level contrasts with the low abundance of monocistronic *gfp* mRNA in the Di-L*atpH*^Zm^ line, showing again that PPR10 translational activation is largely independent of its stabilization of monocistronic RNA.

Upon inserting *rbcL* gene 3’ UTR (T*rbcL*) downstream of *aadA*, the abundance of monocistronic *aadA* transcript dramatically increased and is comparable in amount to the dicistronic mRNA from all four constructs (**upper panel,** **Fig. 3E**). This result indicates T*rbcL* efficiently protects mRNA from degradation in the 3’ end as expected (Stern and Gruissem, 1987). The relative quantities of monocistronic *gfp* to the dicistronic mRNA remained consistent with those without T*rbcL* (**lower panel,** **Fig. 3E, 3A**). The radical difference of monocistronic *aadA* and *gfp* mRNA suggests a lower degree of stabilization of RNA 5’-termini by PPR10 compared to protection of 3’-termini by the stem-loop. The GFP level in the leaves of Di-T*rbcL*+ L*atpH*^Nt^ and Di-T*rbcL*+ L*atpH*^Zm^ dropped to 13% and 20% of TSP (**Fig. 3G**), hypothetically as a result of the synergistic effect of T*rbcL* induced partial RNA cleavage and inefficient transcription termination (compare **Fig. 3G** and **Fig. 3C**) (Stern and Gruissem, 1987). The GG and AA mutations in the maize PPR10 binding site caused a dramatic reduction in GFP accumulation (**Fig. 3G**) as shown previously (Rojas et al., 2019). GFP fluorescence under UV light is consistent with the protein gel data (**Fig. 3D, 3H**). A growth defect was once again observed in the Di-L*atpH*^Nt^, Di-L*atpH*^Zm^, Di-T*rbcL*+ L*atpH*^Nt^ and Di-T*rbcL*+ L*atpH*^Zm^ lines, which can be attributed to the GFP induced metabolic burden and titration of native PPR10 (**Supplemental Fig. 2**).

### Inclusion of tRNA (*trnP*) triggers complete RNA processing, destabilizing *gfp* mRNA at the 5’ UTR

The concept of the IEE has been put forth as a means to facilitate the processing of polycistronic mRNAs and thereby increase the yield of translatable monocistronic mRNAs. On the other hand, there is evidence that the translation activation activity of helical repeat translational activators (such as PPR10) is independent of their stabilization of monocistronic mRNA (see above). To further probe the impact of mRNA cleavage on transgene expression within a multigene operon, we introduced a tRNA *cis*-element between T*rbcL* and L*atpH*^Nt^ in construct Di-T*rbcL*+trnP+L*atpH*^Nt^ (**pAI9,** **Fig. 1C**) (LaManna et al., 2023). Only the monocistronic *aadA* was readily detectable on RNA gel blots indicating efficient cleavage of dicistronic mRNA (**upper panel,** **Fig. 4A**). However, the amount of monocistronic *gfp* mRNA was exceedingly low and became visible only after the film was over-exposed (**lower panel,** **Fig. 4A**). Intriguingly, the abundance of this transcript was notably less than that of the processed monocistronic *gfp* mRNA generated in the Di-L*atpH*^Nt^ and Di-T*rbcL*+L*atpH*^Nt^ plants (**lower panel,** **Fig. 3A, 3E**). This suggests that the RNA cleavage adjacent to the tRNA and subsequent 5’-->3’ RNA degradation is so rapid that PPR10 does not have time to bind and protect the RNA. Quantification via Western blot revealed merely ∼0.4% GFP TSP content in the leaves (**Fig. 4B**, 4C, **Table 1**).

**Figure 4.**
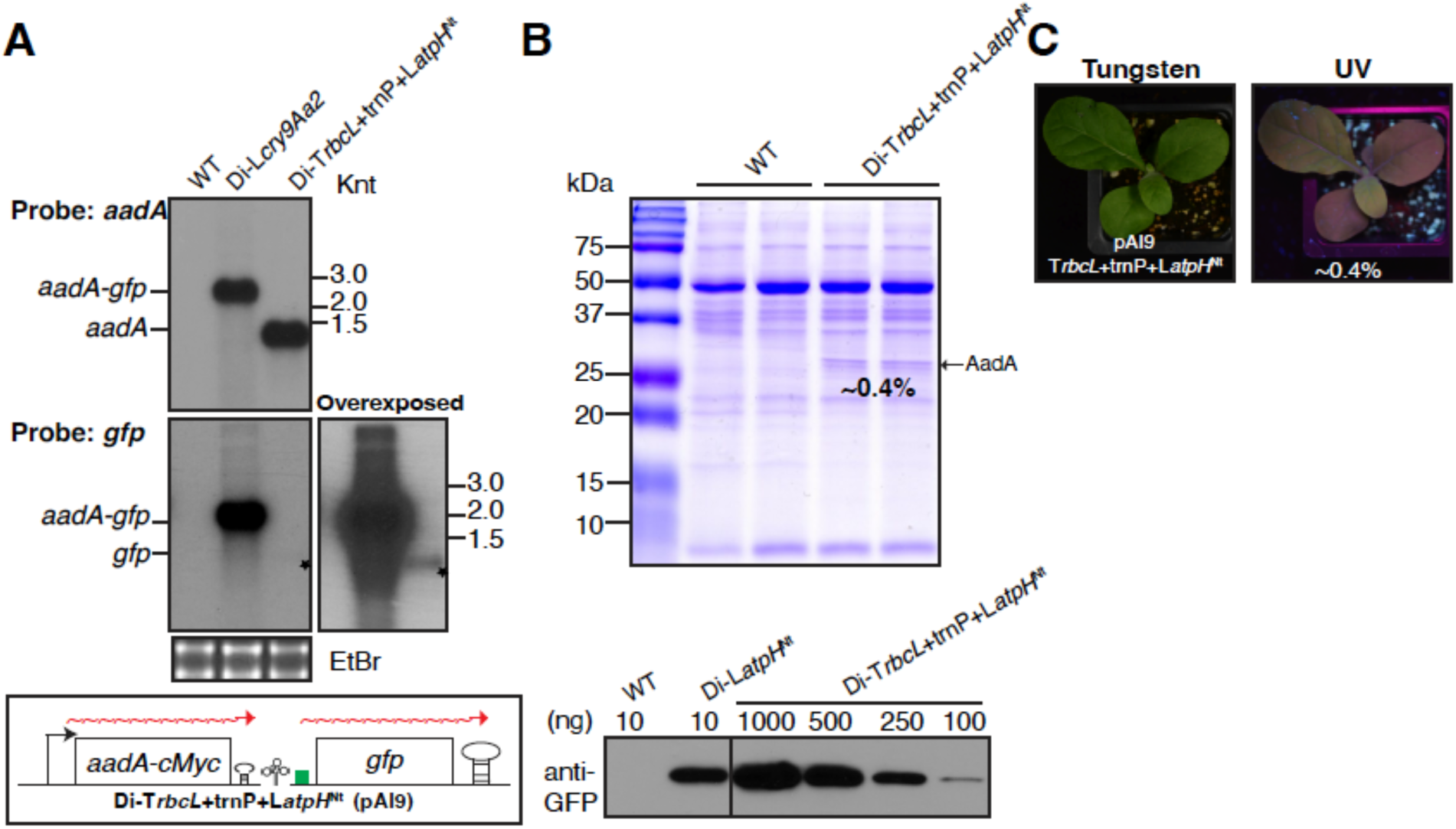
tRNA *cis-*element triggers mRNA cleavage and a strong decrease in transgene expression. **A)** RNA gel blot hybridizations showing only monocistronic *aadA* or *gfp* (asterisks) mRNA accumulation. Knt, Kilonucleotide. **B)** Protein gel blot (upper panel) and Western blot (lower panel) to measure GFP abundance in Di-T*rbcL*+trnP+L*atpH*^Nt^ leaves, using Di-L*atpH*^Nt^ (∼16% of TSP) as the reference. Vertical line separates nonadjacent lanes from the same blot. GFP accumulates at ∼0.4% of TSP (not visible in Coomassie stained gel). **C)** Weak GFP fluorescence is visualized under UV light.

As expected, based on the low GFP expression programmed by either mutating the PPR10 binding site (L*atpH*^GG^) or inserting a tRNA upstream of the *atpH* 5’ UTR, monocistronic *gfp* mRNA and GFP abundance were further diminished when these two features were combined in the same construct (**pAI10,** **Fig. 1C**). The monocistronic *gfp* became barely detectable and GFP accumulated to only ∼0.02% of TSP (**Di-T*rbcL*+trnP+L*atpH*^GG^,** **Fig. 5**).

**Figure 5.**
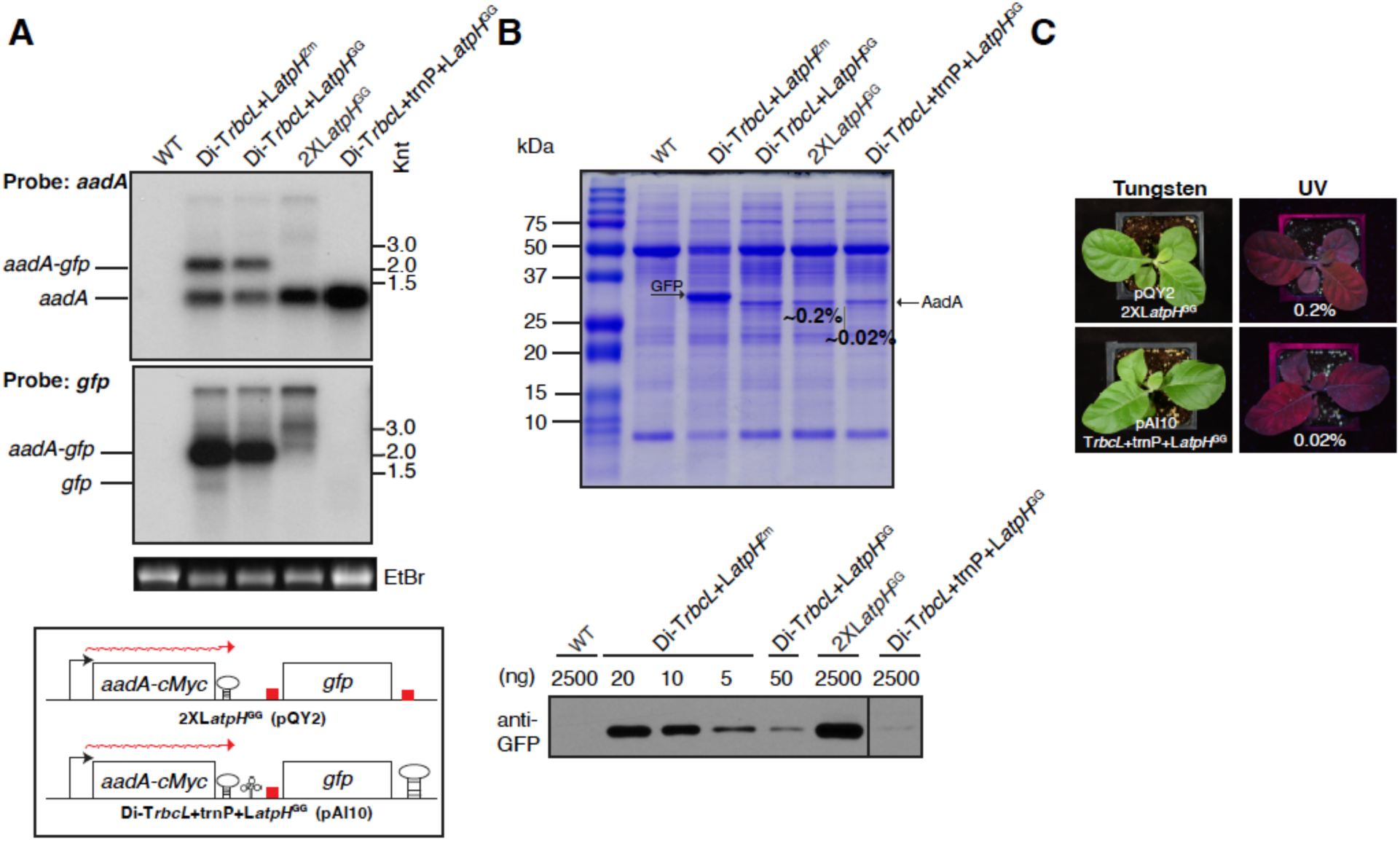
Combination of tRNA *cis-*element with *atpH*^GG^ 5’ UTR or usage of two *atpH*^GG^ 5’ UTRs reduces transgene expression. **A)** RNA gel blot hybridizations showing only monocistronic *aadA* mRNA accumulation. Knt, Kilonucleotide. **B)** Protein gel blot (upper panel) and Western blot (lower panel) to measure GFP abundance in Di-T*rbcL*+tRNA+L*atpH*^GG^ and 2XL*atpH*^GG^ leaves, using total soluble leaf extract from Di-T*rbcL*-L*atpH*^Nt^ (∼15% of TSP) as the reference. Vertical line separates nonadjacent lanes from the same blot. **C)** GFP fluorescence is almost undetectable under UV light.

### Replacement of T*psbA* with L*atpH*^GG^ at the 3’ UTR reduces GFP accumulation and yields readthrough transcripts

As an alternative approach to down regulate gene expression by mediating RNA stability, we replaced T*psbA* with another copy of L*atpH*^GG^ (**pQY2,** **Fig. 1C**). Indeed, only monocistronic *aadA* accumulated, while a well-defined monocistronic *gfp* transcript is absent (**2X L*atpH*^GG^,** **Fig. 5A**). However, unlike the above scenario in which a tRNA was incorporated in the intergenic region, longer readthrough transcripts were detected, which presumably accounts for the greater output of GFP (0.2% GFP out of TSP) (**Fig. 5A, 5B**).

### Titration of endogenous PPR10 by transgenic mRNA harboring the tobacco and maize PPR10 binding site

It was reported previously that inclusion of multiple binding sites for HCF107 in plastid transgenes leads to defects in the stabilization of the transcripts that are stabilized by HCF107, likely via depletion of HCF107 (Legen et al., 2018). When the transplastomic plants described here were grown under heterotrophic conditions in medium containing sucrose for 7 days, cotyledons displayed a range of pale green colors (**Fig. 6A**). All the transplastomic lines harboring the L*atpH*^Nt^ construct displayed a severe pale green phenotype except pAI9, which contained the tRNA *cis*-element. Seedlings with the L*atpH*^Zm^ construct showed a subtle pale green phenotype. Cotyledons of seedlings with L*atpH*^GG^ or L*atpH*^AA^ were all indistinguishable from wild type. The pale green phenotypes became less severe in true leaves among all the lines (**Fig. 6B**). Quantification of the chlorophyll level confirmed those observations (**Fig. 6C, 6D**). The pigment deficiency is clearly not the result of depletion of resources for GFP production, because the maize 5’ UTR lines harbor more GFP than the tobacco 5’ UTR lines. Therefore, it is very likely that the ectopic PPR10 binding site reduces the effective concentration of PPR10. The absence of the pale green phenotype in plants with the tRNA cis-element is presumably related to the fact that monocistronic *gfp* mRNA does not accumulate enough to compete with the native *atpH* mRNA. In addition, when the seeds were germinated and grown in soil in green house conditions, plants displayed similar pigment deficiency patterns in the young emerging leaves as those (L*atpH*^Nt^ and L*atpH*^Zm^) observed in the cotyledons, but the phenotype disappeared gradually as the leaves matured (**Supplemental Fig. 2**). This would argue for the use of heterologous and/or engineered PPR10 proteins and cognate binding sites in chloroplast engineering when high level recombinant protein expression is required, avoiding interference with the native plastid expression system (Rojas et al., 2019; Yu et al., 2019)

**Figure 6.**
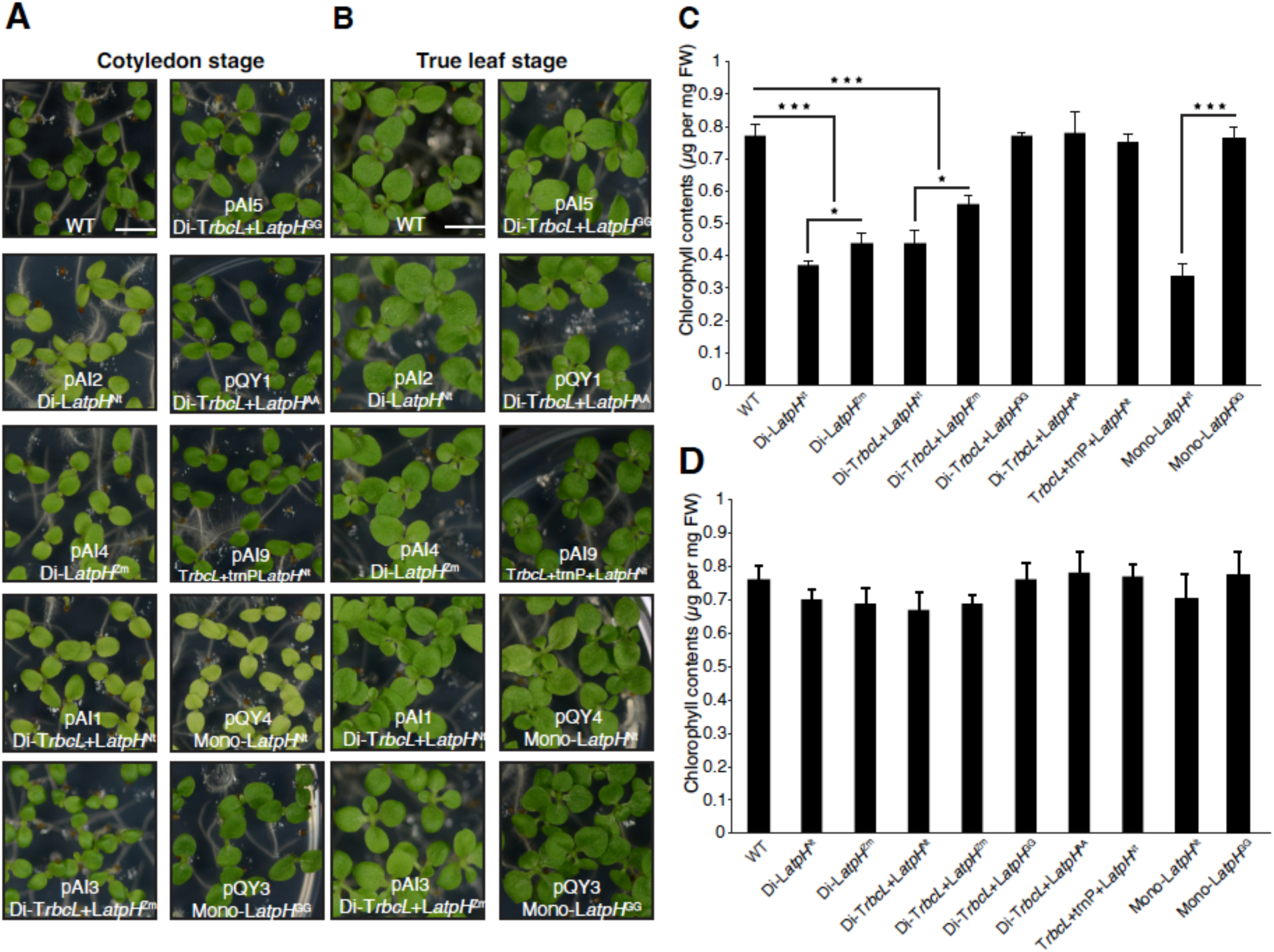
Pigment deficiency in transplastomic plants expressing a transgene with the wild type tobacco and maize PPR10 binding site suggests titration of native PPR10 in cotyledons. **A)** Seedlings with two cotyledons grown in sterile culture on sucrose-containing medium. Note reduced pigmentation in the pAI1, pAI2, pAI4 and pQY4 transplastomic lines, which have a wild-type tobacco or maize PPR10 binding site in the transgene 5’ UTR. **B)** Seedlings with a pair of true leaves grown in sterile culture on sucrose-containing medium. Note uniform green pigmentation in leaves. **C)** Quantification of the chlorophyll content in cotyledons shown in Fig. 6a. **D)** Quantification of the chlorophyll content in leaves shown in Fig. 6b. Differences between the lines were analyzed using Student’s *t*-test. *P<0.05; ***P<0.001, mean±SD, n=5.

## DISCUSSION

This study represents an *in vivo* investigation comparing protein output in tobacco chloroplasts driven by *atpH* 5’ UTR-containing PPR10 binding site variants and the incorporation of a tRNA *cis*-element, aiming to achieve a large dynamic range of plastid transgene expression. These *cis*-elements were primarily tested in a dicistronic context, a configuration which could be positioned in a polycistronic unit to balance the expression of multiple genes.

### Robust mRNA translation from the endogenous tobacco *atpH* 5’ UTR

In a previous study, several PPR footprints, including the tobacco PPR10 binding site that we revisited here, were integrated into a dicistronic reporter construct in tobacco chloroplasts to investigate their impact on plastid transgene expression (Legen et al., 2018). Our L*atpH*^Nt^ constructs yielded much high levels of GFP (∼15% TSP) than those reported in the earlier study. The difference in transgene expression level may result in part from use of different transgene integration sites. Whereas we target the *trnV-3’ rps12* region within an inverted repeat region (IRR) with two transgene copies per ptDNA, the previous study targeted the transgene to a single copy *trnG*-*trnfM* region. Additionally, the 97nt L*atpH*^Nt^ we tested here retained native sequence upstream of the PPR10 binding site, whereas the previous study did not (Legen et al., 2018). These results suggest that sequences upstream of the PPPR10 binding site might play an important role in expression of the downstream gene by optimizing accessibility of the PPR10 binding site.

### Processing of dicistronic mRNAs with L*atpH*^Nt^ and L*atpH*^Zm^ intergenic regions

Most of the dicistronic *aadA*-*gfp* operon constructs in the study share common elements, including the promoter-leader (P*rrn*L*atpB*) and the 3’ UTR (T*psbA*). Notably, L*atpH*^Nt^ generates roughly four times more monocistronic *gfp* mRNA than L*atpH*^Zm^ (**Fig. 3A**), indicating either that the tobacco PPR10 more efficiently stabilizes its cognate binding site or that sequences upstream of the binding site in the tobacco UTR are more susceptible to endonucleolytic cleavage, as cleavage is necessary to produce the substrate for 5’ exonucleolytic processing. Upon the insertion of the *rbcL* 3’ UTR (T*rbcL*) downstream of *aadA*, monocistronic *aadA* transcript accumulated at a level comparable to that of the dicistronic mRNA (**Fig. 3E**), In contrast, very little monocistronic *aadA* accumulated in the absence of T*rbcL* (**Fig. 3A** **vs.** **Fig. 3E**). This suggests that the PPR10 binding site is less effective than T*rbcL* at stabilizing this 3’ end or that there are endonuclease sensitive sites in the *atpH* 5’ UTR upstream of the PPR10 binding site that lead to the degradation of *aadA* sequences in the absence of the stabilizing T*rbcL* element. The abundance of monocistronic *gfp* mRNA remained consistently low, regardless of the presence of T*rbcL*, suggesting that tobacco PPR10 is insufficient to effectively protect the RNA downstream.

### Translation of *gfp* in dicistronic mRNAs with L*atpH*^Nt^ and L*atpH*^Zm^ intergenic regions is not coupled to abundance of processed monocistronic *gfp* mRNA

The maize *atpH* 5’ UTR produced higher output of GFP than the tobacco *atpH* 5’ UTR in both the presence and absence of an upstream T*rbcL* element (**Fig. 3C and 3G**). This finding is noteworthy for two reasons. First, tobacco PPR10 presumably binds with higher affinity to its cognate site than it binds the maize site, which harbors a single nucleotide mismatch. Furthermore, more processed monocistronic *gfp* mRNA accumulated in the L*atpH*^Nt^ lines than in the L*atpH*^Zm^ lines. The fact that the L*atpH*^Zm^ line produces more abundant GFP despite the absence of monocistronic RNA provides strong evidence that translational activation by PPR10 is not solely a result of its stabilization of monocistronic mRNA. It is likely that the same principle applies to other helical repeat activating proteins in chloroplasts, including HCF107, the protein responsible for the activating effects of the IEE element.

We had anticipated that the native tobacco *atpH* 5’ UTR would yield more GFP expression than the heterologous maize *atpH* 5’ UTR, but this was not the case. One hypothesis to explain this counterintuitive observation is that the 3’ boundary of the PPR10 footprint is one nucleotide closer to the 5’ boundary of the footprint of the initiating ribosome in the L*atpH*^Nt^ UTR than in the L*atpH*^Zm^ UTR (**Fig. 1A**). It is conceivable that this proximity places PPR10 so close to the ribosome binding region that it sterically interferes with ribosome binding. Alternatively, the greater titration of endogenous PPR10 by the native tobacco *atpH* 5’ UTR (**Fig. 6**) may lead to a general suppression of plastid translation in chloroplasts due to disrupted ATP synthesis. In any case, these results highlight the utility of employing *cis*-elements that do not precisely match those of the host species.

### Reducing transgene expression by regulating translation efficiency and mRNA stability

The accumulation of GFP levels could be significantly reduced at the posttranscriptional level using two approaches: by regulating RNA translation efficiency and RNA stability. As previously demonstrated (Rojas et al., 2019), the incorporation of point mutations that strongly reduce the interaction between the binding site and PPR10 protein substantially diminished translational activation. Including the GG or AA variants in the dicistronic T*rbcL* plants (pAI5, pQY1) resulted in a reduction of GFP accumulation to 2% and 3% respectively.

The second approach involved a reduction in mRNA abundance by introducing an RNA cleavage *cis-*element at the 5’ end of the target gene. Initially, the motivation behind investigating the effect of inserting a tRNA in the intercistronic region was to enhance the production of processed monocistronic mRNA, often assumed to be more translationally active than polycistronic mRNA (Zhou et al., 2007) However, monocistronic *gfp* mRNA did not accumulate despite the presence of a PPR10 binding site between the tRNA and the *gfp* open reading frame. This suggests that the tRNA processing activity that exposes an unprotected 5’ end upstream of the PPR10 binding site occurs so rapidly following transcription that the downstream RNA is degraded before PPR10 has the opportunity to bind. The GFP expression level dropped from 15% TSP (**pAI1,** **Fig. 3E**, 3G, **Table 1**) to roughly 0.4% TSP (**pAI9,** **Fig. 4**, **Table 1**). When the L*atpH*^Nt^ is replaced by L*atpH*^GG^, GFP expression further decreases to ∼0.02% (**pAI10,** **Fig. 5**, **Table 1**). Furthermore, substituting the T*psbA* terminator with an additional copy of L*atpH*^GG^ at the 3’ end led to significant degradation of the dicistronic mRNA, but did not impede longer transcription readthrough as tRNA does (**pQY2,** **Fig. 5A**).

### Chloroplast engineering applications require stoichiometry control over multiple transgenes

The unique attributes of the chloroplast genetic system make it highly suitable for synthetic biology applications to manipulate traits of agronomic relevance, such as pest resistance (Zhang et al., 2017; Bally et al., 2018) and photosynthesis (Lin et al., 2014; Long et al., 2018; Martin-Avila et al., 2020; Chen et al., 2023). Many of these applications necessitate precise control over the relative expression of multiple proteins to produce multimeric protein assemblies, multi-component signaling systems or muti-enzyme biosynthesis pathways. For example, the carboxysome, a proteinaceous organelle encapsulating Rubisco and carbonic anhydrase (CA), assimilates carbon within a selectively permeable protein shell in cyanobacteria. Its stoichiometric composition plays an essential role in determining shell integrity, permeability and catalytic performance (Liu et al., 2021). Quantification of the protein components in *H. neapolitanus* carboxysomes structure revealed an extreme stoichiometry with the highest and lowest functional unit per carboxysome at a ratio of 500: 1 (Sun et al., 2022). Additionally, achieving high titers synthesis of natural products typically requires the coordinated expression of multiple genes at an appropriate stoichiometry to minimize the accumulation of toxic intermediates (Paddon and Keasling, 2014). The tRNA *cis*-element characterized here can be valuable to dramatically reduce protein accumulation from an ORF that is part of an operon transcribing a single primary transcript. These PPR10 binding site variants and tRNA processing elements can then be combined with the inducible engineered PPR10 variants we reported previously (Rojas et al., 2019; Yu et al., 2019) or with inducible systems imported from heterologous systems, such as the IPTG-inducible expression system based on the Lac repressor (Muhlbauer and Koop, 2005) or theophylline-responsive riboswitch (Verhounig et al., 2010; Emadpour et al., 2015).

## Methods

### Plant Material and Growth Conditions

*Nicotiana tabacum* cv. Petit Havana (tobacco) plants were maintained aseptically in Magenta Boxes on RM medium containing Murashige and Skoog salts and 3% sucrose (pH=5.8) to generate leaf material for plastid transformation experiments. The RMOP medium used for tobacco plastid transformation is based on RM medium supplemented with benzyladenine (1 mg/L), naphthaleneacetic acid (0.1 mg/L) and thiamine (1 mg/L) (Lutz et al., 2006). Plants were grown in a standard growth chamber illuminated with cool-white fluorescent tubers for 12 h at 28°C during the day and 28°C during the night.

### Plastid transgene constructs and transformation

The monocistronic construct pQY3 (Mono-*atpH*^GG^) and the dicistronic constructs pAI5 (MK482728) and pQY1 (MK482731) were reported previously (Rojas 2019). The plasmid pQY4 is derived from pQY3 by replacing the *atpH*^GG^ with *atpH*^Nt^, a 96-nt segment of the upstream of *atpH* (Supplementary Table S1).

The plasmid pMRR13 (L*Cry9Aa2*) is the progenitor of the dicistronic constructs containing *atpH* 5’ UTR variants and tRNA. Plasmid pMRR13 has been described (Yu et al., 2020). The DNA sequence of intergenic region between the *aadA* stop codon and the *Nhe*I site at the *gfp* N-terminus is included in Supplementary Table S1. Plasmid pQY2 was obtained by replacing T*psbA* with L*atpH*^GG^ in plasmid pAI5. DNA sequence from the *aadA* stop codon till *Hin*dIII site is shown in Supplemental Table S1. Synthetic DNA has been purchased from BlueHeron Biotechnology, Bothell, WA.

Plastid transformation was carried out as described (Lutz et al., 2006; Maliga et al., 2021). Plasmid DNA for plastid transformation was prepared using the QIAGEN Plasmid Maxi Kit (Cat. No 12162; Qiagen inc., Valencia, CA). Leaves were placed abaxial side up on the RMOP medium. The transforming DNA was coated on the surface of gold particles (0.6 μm, BIO-RAD, Hercules, CA) and introduced into the leaf chloroplasts using Du Pont PDS1000He Biolistic gun (DuPont, Wilmington, DE). Two days after bombardment, the leaf sections were transferred to RMOP medium containing 500 mg/L spectinomycin dihydrochloride pentahydrate (Duchefa Biochemie B.V., Haarlem, Netherlands, Prod. No. S0188.1000). The spectinomycin resistant clones were identified as green shoots 6 to 12 weeks after bombardment. Leaves from the regenerated shoots were transferred onto the same RMOP medium for purification. The uniform population of transformed plastid genome copies was confirmed by DNA gel blot analysis (Supplementary Fig. 1).

### DNA gel blot analysis

Total leaf DNA was prepared by the cetyltrimethylammonium bromide (CTAB) protocol (Yu et al., 2017). Total cellular DNA was digested with the *Bam*HI restriction enzyme. 2 μg DNA loaded per lane was separated by electrophoresis on 1% agarose gels and transferred to Hybond-N membranes (GE Healthcare, Chicago, IL) using capillary flow transfer method. The membrane was hybridized to random-primed ^32^P labeled probe (GE, Healthcare, Chicago, IL) in a modified Church hybridization buffer (0.5M phosphate buffer, pH 7.2, 10mM EDTA, 7% SDS (Church and Gilbert, 1984) overnight at 65^0^C). DNA probe was prepared with *Apa*I/*Bam*HI ptDNA fragment encoding part of the 16S rRNA gene.

### RNA gel blot analysis

Total cellular RNA was isolated from leaves frozen in liquid nitrogen using TRIzol (Thermo Fisher Scientific, Waltham, MA), following the manufacturer’s protocol. 2 μg RNA was electrophoresed on 1.5% agarose/formaldehyde gels, which was transferred to the Hybond-N membranes (GE Healthcare, Chicago, IL) using capillary flow transfer method. Radioactive probes were prepared by random-primed ^32^P labeling (GE Healthcare, Chicago, IL) of gel-purified DNA fragments of the following genes: *aadA*, 0.8-kb *Nco*I-*Xba*I fragment from plasmid pHC1 (Carrer et al., 1991); *gfp*, fragment amplified from *gfp* coding region using primers *gfp*-forward p1(5’-TTTTCTGTCAGTGGAGAGGGTG-3’) and *gfp*-reverse p2 (5’-CCCAGCAGCTGTTACAAACT-3’). Hybridazation results were imaged with a Typhoon RGB scanner (GE Life Sciences, Chicago, IL) and quantified with ImageJ software (version 2.0.0-rc-38/1.50b).

### Protein extraction and Western blot analysis

We followed the protocol reported previously (Kuroda and Maliga, 2001). About 200 mg leaf was homogenized in 200 μl buffer containing 50 mM Hepes/KOH (pH7.5), 10mM potassium acetate, 5mM magnesium acetate, 1mM EDTA, 1mM DTT, 2mM PMSF and 1% β-mercaptoethanol. Insoluble material was removed by centrifugation. Protein concentrations were determined by Bradford protein assay reagent kit (BIO-RAD, Hercules, CA). Visible GFP bands in the Comassie Blue Stained protein gels were quantified with 1D-Multi lane densitometry (Alpha Innotech Alphaimager 2200, San Leandro, CA). GFP in tissues containing very low GFP levels was quantified by Western analyses using transplastomic plants with known amounts of GFP as reference. Each value is an average of measurements of three biological replicates. GFP was detected with a monoclonal mouse anti-GFP antibody (JL-8, Takara Bio USA, San Jose, CA) in Western blot analyses.

### Chlorophyll quantification

Seeds were sterilized with the treatment of 0.85% (w/v) sodium hypochlorite (10X diluted 8.5% (w/v) commercial bleach) in a 1.5mL Eppendorf tube for 3min. The seeds were then washed three times with sterile distilled water. Seeds were germinated on RM medium in deep petri dish (20 mm high and 10 cm in diameter) in the growth chamber illuminated with cool-white fluorescent tubers for 12 h at 28°C during the day and 28°C during the night. 6-8 fresh seedlings with two cotyledons or a pair of true leaves from each line were weighed and homogenized with 1 mL methanol. The homogenized mixture was incubated in the dark for 20 min and centrifuged at 800 rpm for 15 min. A spectrophotometer was used to measure A665 and A652 of the supernatant and Chlorophyll-a, Chlorophyll-b, and Chlorophyll-a+b were calculated using following equations: Chlorophyll-a=18.22\**A*^665.2^-9.55\**A*^652.0^; Chlorophyll-b=33.78\**A*^652.0^-14.96\**A*^665.2^; Chlorophyll-a+b=24.23\**A*^652.0^+3.26\**A*^665.2^ (Porra et al., 1989)

## Data availability

The data supporting the findings of this study are available within the paper and its supplementary information files.

## Acknowledgements

We are very grateful to Mary Galli for critical reading, editing and useful comments and discussions on the manuscript. This research was supported by a grant from USDA NIFA Foundational Program Award Number 2014-67013-21600 to A.B. and P.M. and the National Science Foundation Grant NSF MCB 1716102 to P.M. and Kerry A. Lutz.

## Author Contributions

P.M., T.T.H. and A.B. conceived the project and designed the strategy. T.T.H. supervised undergraduate Alexander Ioannou, who constructed most of plastid transformation vectors and obtained transplastomic line pAI3. Q.Y. performed plastid transformation, obtained transplastomic plants, characterized gene expression and interpreted data. P.M., Q.Y. and A.B. wrote the manuscript.

## FIGURE LEGENDS

**Supplemental Figure S1.** DNA gel blot confirms the uniform transformation of the plastid genomes. *Bam*HI digested total cellular DNA was probed with the 2.1-kb 16S rRNA *Apa*I/*BamH*I probe. Knt refers to Kilonucleotide.

**Supplemental Figure S2.** Impaired growth of transplastomic plants expressing the *gfp* reporter gene with the tobacco and maize *atpH* 5’ UTR.

**Supplemental Figure S3.** Annotated sequence file of the vectors in the study.

**Supplemental Table S1.** DNA sequence of vectors in the study.

